# More than fruity scents: floral biology, scent and spectral reflectance of Annonaceae species

**DOI:** 10.1101/2024.09.16.613362

**Authors:** Ming-Fai Liu, Junhao Chen, Chun-Chiu Pang, Tanya Scharaschkin, Richard M. K. Saunders

**Author notes:** Correspondence 1: Ming-Fai Liu, Flora Conservation Department, Kadoorie Farm & Botanic Garden, Lam Kam Road, Lam Tsuen, Hong Kong;, 2: Richard M.K. Saunders, Area of Ecology & Biodiversity, School of Biological Sciences, The University of Hong Kong, Pokfulam Road, Hong Kong.

## Abstract

**Premise:** The family Annonaceae possesses a broad array of floral phenotypes and pollination specialisations, and are important in the plant-pollinator interactions of tropical rainforests. Although there has been considerable effort to assess their interactions with pollinators, attempts to characterise their visual and olfactory communication channels are scarce.

**Methods:** Here, we investigated the pollination biology of 12 Annonaceae species from five genera, viz. *Meiogyne*, *Monoon*, *Polyalthia*, *Pseuduvaria*, and *Uvaria*. Furthermore, their floral colour was characterised by reflectance spectroscopy and floral odour chemistry was assessed using gas chromatography-mass spectrometry. Floral scent was further compared across the whole family using non-metric dimensional scaling plots to identify specialisation in floral odour.

**Results:** The *Meiogyne* species are likely pollinated by small beetles; the *Polyalthia* and *Pseuduvaria* species are likely pollinated by beetles and flies; and the *Uvaria* species is likely pollinated by beetles and bees. Flowers of most species are UV non-reflective, and have various spectral reflectance profile across the remaining visible spectra. Multiple species produce floral odour resembling ripe fruits. The flowers of *Meiogyne* species and *Polyalthia xanthocarpa* emitted mostly branched-chain esters, while flowers of *Uvaria* released mainly straight-chain esters. The *Pseuduvaria* species instead emitted scent reminiscent of rotten fruits, largely consisting of 2,3-butanediol. The inner petal corrugation in *Meiogyne* functions as a food reward, and the inner petal growth serves as a nectary gland for *Pseuduvaria*.

**Conclusions:** Our study identifies the visual and olfactory cues of multiple Annonaceae species and provides insights into how Annonaceae flowers attract different guilds of pollinators.

## INTRODUCTION

The pantropical family Annonaceae is an early-divergent angiosperm lineage of trees and lianas, comprising ca. 2500 species from 110 genera (Couvreur et al., 2019). Annonaceae species are one of the major components of tropical lowland to lower montane rainforests (Sosef et al., 2017). Almost all Annonaceae species possess a basic floral architecture of one whorl of three sepals, two whorls of three petals, numerous spirally arranged stamens and often free carpels (van Heusden, 1992). These flowers are largely pollinated by small beetle pollinators from Curculionidae, Nitidulidae, Staphylinidae and Chrysomelidae (Gottsberger, 2012; Saunders, 2012). Because beetle pollinators spend considerable time in the flowers and can often inflict damage to floral tissues, Annonaceae flowers have evolved protective structures like fleshy corolla and shield-like stamen connectives (Gottsberger, 1999). These adaptations are particularly pronounced in Neotropical species that are pollinated by large scarab beetles from Cetoniinae, Dynastinae, Rutelinae and Trichiinae (Gottsberger, 1999). While small beetles pollinate the majority of Annonaceae species, some are reported to be pollinated by thrips (Gottsberger, 1999) or small flies (Silberbauer-Gottsberger et al., 2003) and more rarely, flowers can be pollinated by bees (Teichert et al., 2009) or cockroaches (Nagamitsu et al., 1997). Pollinators are often rewarded with pollen, stigmatic exudates, floral nectar and thickened nutritious floral tissues (Gottsberger and Webber, 2018; Saunders, 2020). The floral chamber created by the corolla often additionally serves as tryst sites, and more rarely as oviposition sites (Saunders, 2020). Highly specialised pollination systems have been previously identified in Annonaceae, such as floral mimicry of fruits (Goodrich and Jürgens, 2018, and references therein), fermentation substrates (Goodrich et al., 2006), mushrooms (Teichert et al., 2012) and aerial litter (Liu et al., 2024). Other derived pollination strategies, including private channel communication with beetles (Maia et al., 2012) and euglossine bees (Teichert et al., 2009) have also been reported. Among these floral specialisations, mimicry of fruits was proposed to be the ancestral condition for the family (Johnson and Schiestl, 2016), and was likely pervasive, as around half of the recorded species emit fruity floral aromas (Goodrich, 2012).

How flowers achieve specialised pollination is a research interest for pollination ecologists. In particular, visual and olfactory cues have been recognised for their roles in mediating these interactions (Schaefer and Ruxton, 2011). Annonaceae flowers display diverse combinations of colour and odour, which might have enabled the evolution of specialisation in pollinators. Strong floral fragrance is often noticeable in Annonaceae flowers. Based on a review by Goodrich (2012), there appears to be a myriad of floral odour reported in the family based on textual description. Although delineation of the chemical components of floral aromas is rare for Annonaceae, volatiles from over 12 chemical classes have been reported so far (Goodrich, 2012). The type of pollination adaptation seems to be associated with molecules of specific classes. For instance, proposed fruit mimics appear to be associated with aliphatic esters (Jürgens, 2000; Johnson and Schiestl, 2016); flowers mimicking fermenting substates reportedly emit fermentation by-products, including short-chain alcohols and ketones (Goodrich et al., 2006); and monoterpenoids are associated with aerial litter mimics (Liu et al., 2024) and flowers pollinated by Euglossine bees (Teichert et al., 2009).

Floral coloration is also diverse within the family, and is more frequently documented in the literature. Floral colours, including yellow, red, pink, green, orange, white and dark maroon or indigo flowers have been recorded in Annonaceae (van Heusden, 1992). It has been reported that colour can shape the choice decision of one of the major pollinating beetle family Nitidulidae (Döring et al., 2012; Vuts et al., 2022), highlighting the importance of floral visual cues. Surprisingly, virtually no attempt has been made to characterise Annonaceae floral colour based on spectral reflectance with only one exception (Liu et al., 2024). Given the diversity of pollination strategies, floral colour is an overlooked area which can potentially help elucidate how Annonaceae flowers interact with their pollinators.

Previous studies have revealed that Annonaceae species in Australia display a variety of floral adaptations to small beetle, fly or bee pollinators (Silberbauer-Gottsberger et al., 2003). For instance, *Meiogyne* and *Polyalthia* (as *Haplostichanthus*), both generally with yellowish corolla, are reportedly pollinated by small beetles from the family Nitidulidae and Curculionidae, respectively. *Pseuduvaria* is reportedly pollinated by flies, and *Uvaria* is reportedly pollinated by stingless bees. The aim of the current study is to characterise the phenotypic adaptations for various types of pollination systems in Annonaceae. We selected 12 species from the genera *Meiogyne*, *Monoon*, *Polyalthia*, *Pseudvaria* and *Uvaria* in Australia. These systems offer arenas for studying specialised bee pollination and potential floral mimicry of fruit and fermenting substrates. Their floral biology, floral spectral reflectance and floral odour are characterised here, laying a foundation for the future study of floral traits and pollination ecology of specialised pollination systems. Furthermore, magnoliids including Annonaceae preserve many architectural elements of ancestral angiosperm flowers (Sauquet et al., 2017). Characterisation of the pollination strategies employed by early-divergent angiosperms like magnoliids may therefore offer insights into the pollination system of the ancestral flower.

## MATERIALS AND METHODS

### Study species and study sites

We examined three *Meiogyne* Miq. species (*Meiogyne cylindrocarpa* (Burck) Heusden, *Meiogyne trichocarpa* (Jessup) D.C.Thomas & R.M.K. Saunders, *Meiogyne stenopetala* (F.Muell.) Heusden), one *Monoon* Miq. species (*Monoon patinatum* (Jessup) B.Xue & R.M.K.Saunders), two *Polyalthia* Blume species, (*Polyalthia hispida* B.Xue & R.M.K.Saunders, *Polyalthia xanthocarpa* B.Xue & R.M.K.Saunders), four *Pseuduvaria* Miq. species (*Pseuduvaria hylandii* Jessup, *Pseuduvaria glabrescens* (Jessup) Y.C.F.Su & R.M.K.Saunders, *Pseuduvaria mulgraveana* Jessup, *Pseuduvaria villosa* Jessup) and two *Uvaria* species (*Uvaria concava* Teijsm. & Binn. and *Uvaria rufa* Blume) (See Table 1 for detailed information on the studied plants. Observations were conducted in the wild for individuals of *Me. stenopetala* in Mount Tamborine in 2013 (permit number: WITK08297010). All other observations were made on cultivated individuals at various sites: Gardens by the Bay, Singapore, in 2018; Garry and Nada Sankowsky Arboretum, Tolga, Australia, in 2019 and 2023; Cairns Botanic Gardens, Australia, in 2023; and Graham Wood Arboretum, Wondecla, Australia, in 2023 (see Table 1 for details).

**Table 1.**
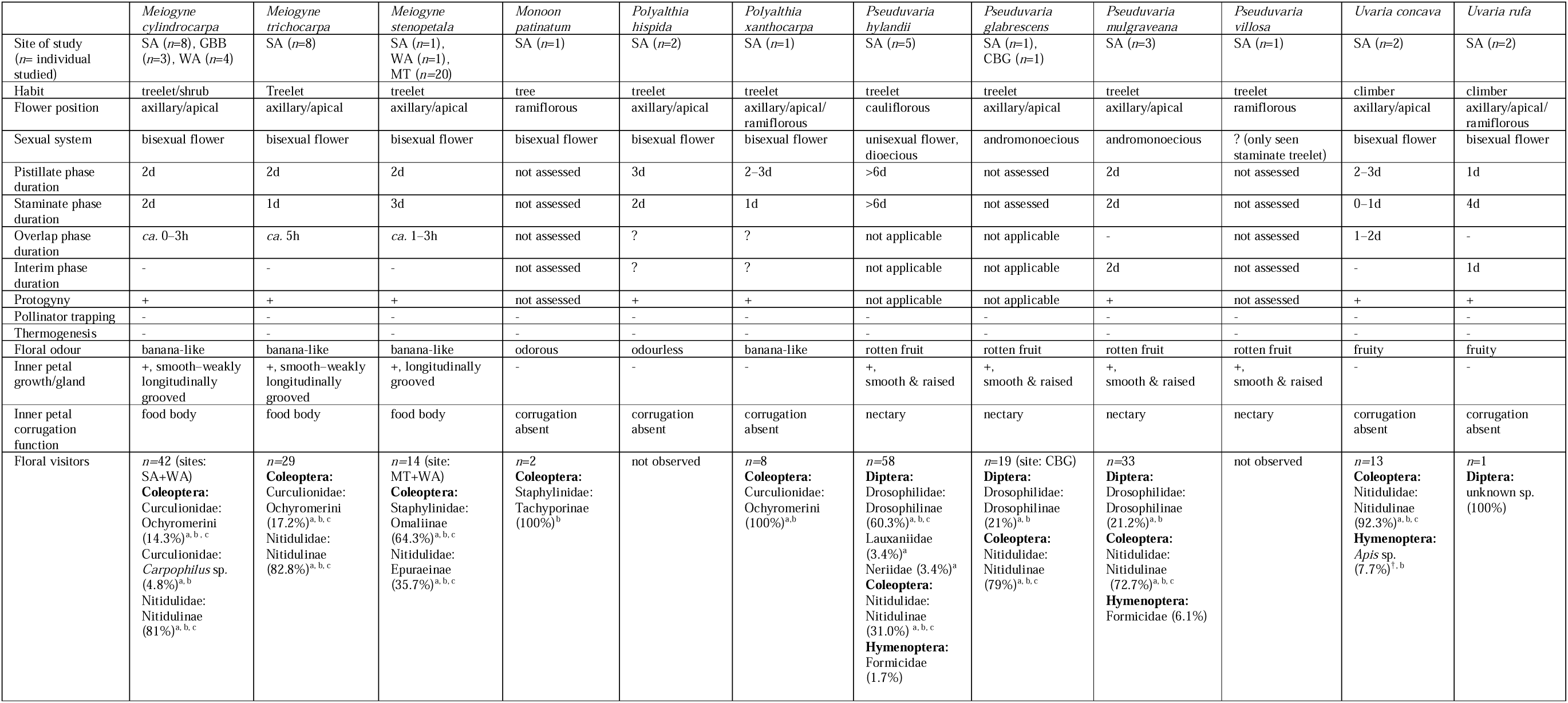
Floral biology of *Meiogyne*, *Monoon*, *Polyalthia*, *Pseuduvaria* and *Uvaria* species. Abbreviations: CBG: Cairns Botanic Gardens, NE Queensland, Australia; GBB: Gardens by the Bay, Singapore; MT: Mt. Tamborine, SE Queensland, Australia; SA: Sankowsky Arboretum, Tolga, NE Queensland, Australia; WA: Graham Wood Arboretum, Wondecla, NE Queensland, Australia; +: present, -: absent, a : presence on both pistillate-phase and staminate-phase flowers, b : contact with reproductive organs, c: pistillate-phase visitors were pollen-laden.

### Phenology

To assess floral ontology, 10 flowers for each species were tagged and observed for seven consecutive days. Observations were made daily to assess the duration of the pistillate and staminate phase. Stigmatic peroxidase activity has been widely used as a gauge for stigma receptivity (Dafni and Maués, 1998). However, the stigmas of some species, especially *Meiogyne* spp., displayed peroxidase activity even during bud stage based on our preliminary assessment with Peroxtesmo KO paper (MacheryNagel, Düren, Germany). The onset of stigmatic receptivity was therefore defined by stigmatic exudate secretion, while the end of stigmatic receptivity was determined by the abscission of stigmas. Staminate activity was similarly determined by anther dehiscence and the abscission of petals. A pistillate phase is defined here as the period that the flower has only the pistillate function, and likewise a staminate phase is defined as the period that the flower only exhibits the staminate function. An overlap phase describes the period that the flower has both the pistillate and staminate functions, while an interim phase refers to the non-receptive period between the pistillate and staminate phase. Floral thermogenesis was assessed using a handheld digital thermometer every two hours throughout the day in the floral chamber, with additional readings ca. 10 cm away from the flower to measure ambient temperature.

### Floral visitors

Floral visitors were recorded during the entire observation period. A representative number of floral visitors were collected and either preserved in 50% isopropanol for identification or frozen at −20°C and silica-dried for inspection of pollen deposition under a dissecting microscope. Probable pollinators were assessed by these criteria: 1) visitation to flowers of both sexes or sexual phases; 2) contact with floral sexual organs; and 3) presence of pollen grains on visitors in pistillate-phase flowers. Since the pistillate phase precedes the staminate phase, the presence of these pollen-laden insects provide evidence for inter-floral transfer of pollen grains.

### Spectral reflectance of the petals

Pollinators often access Annonaceae floral reproductive organs via apical and basal apertures created by the inner petals (Saunders, 2012). For genera with similar colour and morphology between the inner and outer petals, namely *Meiogyne*, *Monoon*, *Polyalthia* and *Uvaria*, petal spectral reflectance was represented by that of the inner petals. The abaxial surface of the inner petals was measured for genera with drooping inner petals, viz. *Meiogyne*, *Monoon* and *Polyalthia*, whereas the adaxial surface of the inner petals were measured for *Uvaria*, which has spreading inner petals. *Pseuduvaria* has varying coloration among different parts of the petals, and therefore three perianth areas were measured: outer petal adaxial surface, inner petal abaxial surface, and inner petal gland. Differences between sexes and sexual phases were not assessed due to logistic constraints. Spectral reflectance of the flowers at 300–700 nm was measured using an Ocean Optics FLAME-S-UV-VIS-ES spectrometer (Dunedin, Florida, USA), a DH-mini UV-VIS-NIR light source and an R200-7-UV-VIS probe positioned at 45°. For each species, individual flowers (n=1–8) selected randomly from one to five biologically distinct individuals were used as replicates.

### Floral odour

To sample the floral volatiles, up to 10 intact flowers were enclosed in a “Toppits” oven bag for each sample. To control for ambient contaminants, an empty bag was likewise sampled. Two slits were made in the headspace bag, with one end connecting to a charcoal cartridge (ORBO 32, Supelco, Pennsylvania, USA), supplying filtered air, and the other connected to a PAS 500 air pump (Spectrex, California, USA). Floral volatiles were collected in a Porapak-Q cartridge (ORBO 1103, 50/80, Supelco, Pennsylvania, USA) at a flow rate of 100 m/min for 4 h. The porapak-Q cartridge was then sealed and transported to the laboratory at The University of Hong Kong. The floral volatiles then were eluted in 1 ml hexane (HPLC-grade, Sigma-Aldrich, Massachusetts, USA). The eluents were then concentrated to 200 µl by nitrogen blowdown. For each species, technical replicates were made. The internal standard toluene was added to each sample at 34.68 ng/µl.

Gas Chromatography-Mass Spectrometry was performed on an Agilent 6890N/5973 gas chromatograph-mass selective detector to characterise floral headspace. For each sample, 1 µl eluent was injected into the inlet at 230 °C for 1 min, and then injected into a DB-WAX column (30 m × 0.25 mm, 0.25 μm thick film; J&W) in splitless mode. Helium was supplied at a flow rate of 1 ml/min as the carrier gas. Oven temperatures were held at 60 °C for 3 min, followed by a ramp up to 250 °C at 10 °C/min, and finally held for 7 min. Peaks were manually integrated and identified tentatively using the Wiley and National Institute of Standards and Technology mass spectral libraries (NIST, Maryland, USA; 2020), with an 85% quality threshold. Compound identities were verified by published standardised retention index values or by co-injecting with commercially available standards. Molecules were quantified with the internal standard toluene (see also Kantsa et al., 2018). The relative abundance of each molecule was calculated. Volatiles recovered in the ambient control sample were excluded in other samples.

To assess the similarity of the floral odour among Annonaceae species, non-metric dimension scaling (NMDS) analysis was performed, expanding on the sampling of a previous family-wide NMDS study (Jürgens, 2009). The floral odour data generated in the current study were supplemented by published records of floral odour in the family (Jürgens, 2000; Goodrich et al., 2006; Goodrich and Raguso, 2009; Teichert et al., 2009, 2012; Gottsberger et al., 2018; Chen et al., 2020; Liu et al., 2024). Unknown molecules were excluded. When stereochemistry is unclear, enantiomers were added together in the matrix. The data were log-transformed and Bray-Curtis dissimilarity index was calculated, which was then used to construct an NMDS plot with the R package vegan. The floral odours were assigned with the type of pollination systems based on the findings of the current study and the literature (Silberbauer-Gottsberger et al., 2003; Jürgens, 2009; Goodrich et al., 2006; Goodrich & Raguso, 2009; Teichert et al., 2009; 2012; Gottsberger et al., 2018; Chen et al., 2020; Liu et al., 2024). Analysis of Similarity (ANOSIM) assesses if similarity between groups are statistically different from within groups.

## RESULTS

The floral biology of the studied species is summarised in Table 1. Most species assessed displayed strong levels of protogyny and none were thermogenic. All *Meiogyne*, *Monoon*, *Polyalthia* and *Uvaria* species assessed here bear hermaphroditic flowers, whereas the *Pseuduvaria* species bear unisexual flowers (*Ps. hylandii* and *Ps. glabrescens*) or a combination of staminate and hermaphroditic flowers (*Ps. mulgraveana*).

### Floral biology and pollinator activities—*Meiogyne* species

The three *Meiogyne* species (Fig. 1a–c) produced yellow flowers with longitudinally grooved basal outgrowths on the adaxial surface of the inner petals (Fig. 1c, Fig. 2a, c) and emitted a banana-like scent. Their pistillate phase lasted two days, while the duration of staminate phase differs among species (1–3 days) (Table 1). They exhibited a short overlap between the pistillate and staminate phases that lasts ca. 0–5 hours. Small beetles were frequently observed visiting the flowers of *Me. cylindrocarpa* (Curculionidae and Nitidulidae), *Me. trichocarpa* (Curculionidae and Nitidulidae), and *Me. stenopetala* (Nitidulidae and Staphylinidae) in both staminate and pistillate phase, during which they contacted the stamens and stigmas. Pollen-laden curculionid beetles from the tribe Ochyromerini and pollen-laden nitidulid beetles from the subfamily nitidulinae observed in pistillate-phase flowers were likely the pollinators of both *Me. cylindrocarpa* and *Me. trichocarpa*. The staphylinid beetles from the subfamily Omaliinae and the nitidulid beetles from the subfamily Epuraeinae were likely the pollinators of *Me. stenopetala*. These floral pollinators consumed pollen grains on the flowers where the beetles also sought shelter and copulated. Stigmas of all three *Meiogyne* species produced a thin film of stigmatic exudate, which became more copious towards the end of the pistillate phase. Consumption of stigmatic exudate was invariably observed in curculionid, nitidulid and staphylinid beetle visitors. Consumption of floral tissues, specifically petals and stigmas, were observed, primarily by curculionid beetles, and less frequently by nitidulid and staphylinid beetles, leaving gnaw marks on the floral tissues of the three *Meiogyne* species. However, consumption of the inner petal corrugation was particularly pronounced compared to the surrounding floral tissues (Fig. 2). Nectar secretion was not observed on the inner petal corrugation of *Me. cylindrocarpa*, *Me. trichocarpa* and *Me. stenopetala*, thus they likely functioned as a food body reward for the floral visitors.

**Fig. 1.**
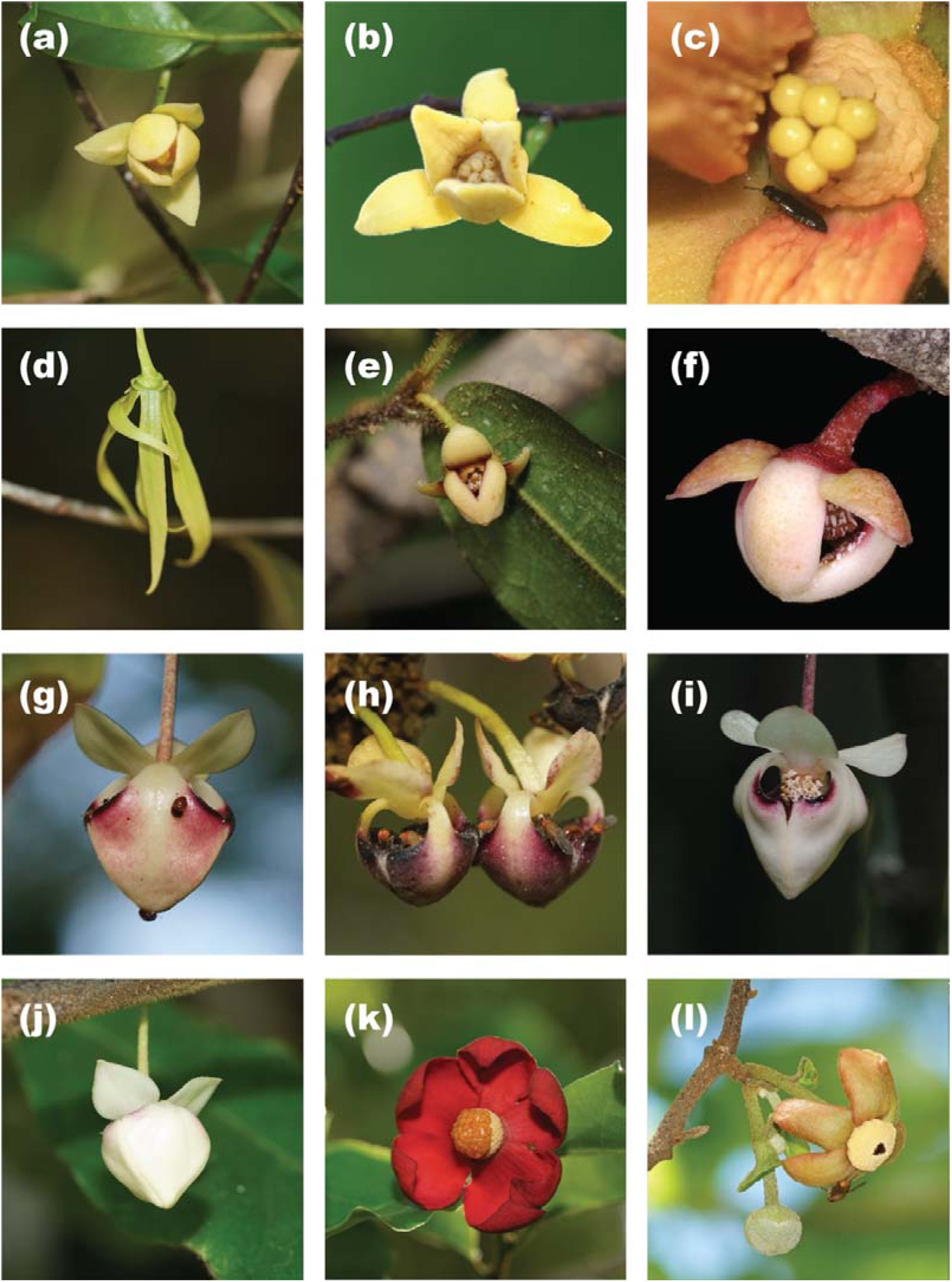
Flowers of Annonaceae species studied. (a) *Meiogyne cylindrocarpa*. (b) *Meiogyne trichocarpa* with curculionid and nitidulid visitors. (c) *Meiogyne stenopetala* with a staphylinid visitor. (d) *Monoon patinatum*. (e) *Polyalthia hispida*. (f) *Polyalthia xanthocarpa.* (g) *Pseuduvaria glabrescens* with nitidulid visitors. (h) *Pseuduvaria hylandii* with drosophilid visitors. (i) *Pseuduvaria mulgraveana*. (j) *Pseuduvaria villosa*. (k) *Uvaria concava*. (l) *Uvaria rufa* at interim phase with a fly visitor. Photo credit: Photo credits: (a, b, d, e, g–l): Ming-Fai Liu; (c): Chun-Chiu Pang; (f): Garry & Nada Sankowsky.

**Fig. 2.**
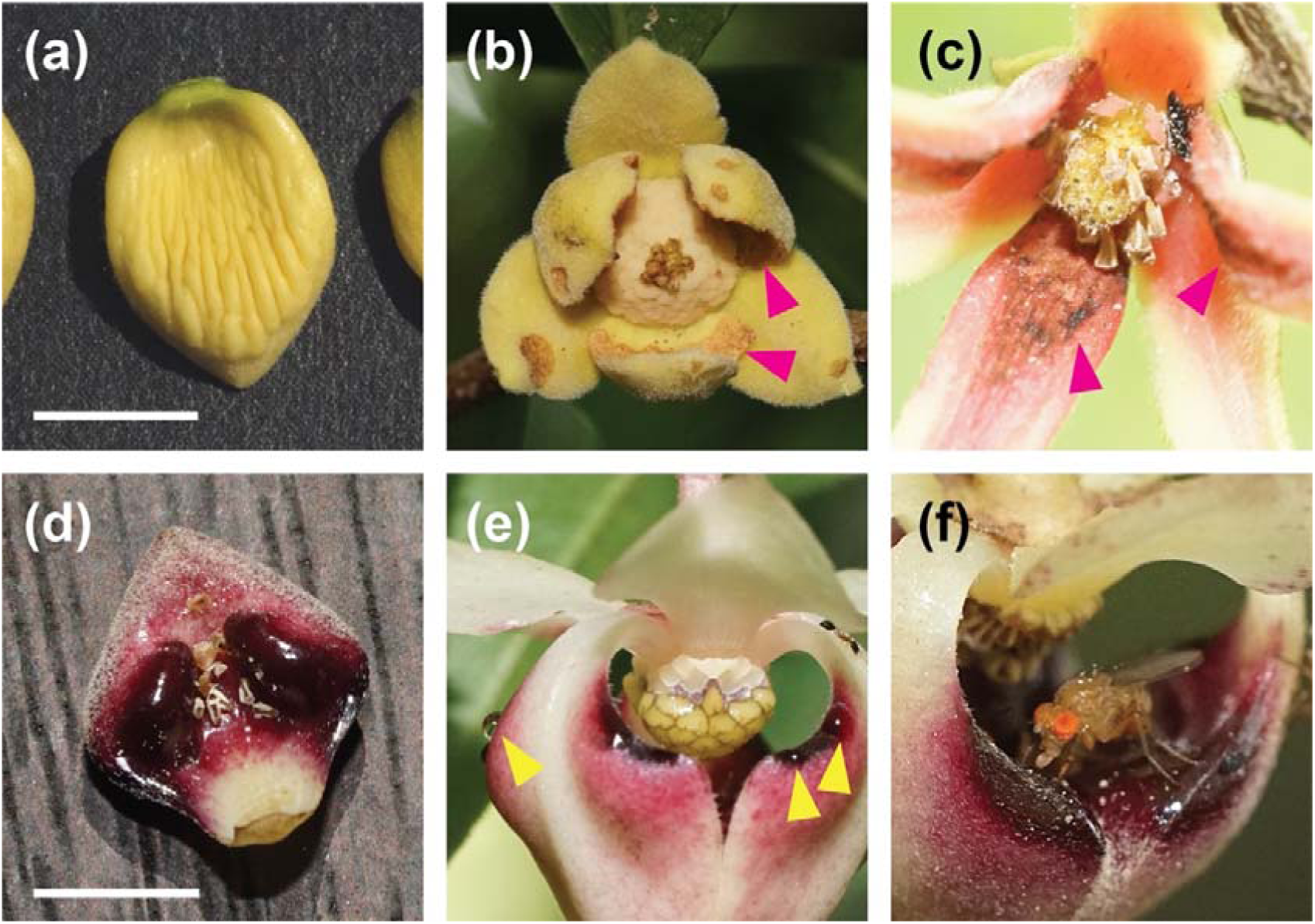
Inner petal outgrowths. (a) Adaxial surface of inner petal of *Meiogyne cylindrocarpa*. (b) Gnaw marks (magenta arrows) created by curculionid beetles on the inner petals of *Meiogyne trichocarpa*. (c) Gnaw marks (magenta arrows) created by staphylinid beetles on the inner petals of *Meiogyne stenopetala*: longitudinal ridges of the basal corrugation have been gnawed away. (d) Pair of nectary glands on the adaxial surface of the inner petal of *Pseuduvaria hylandii*. (e) Nectar secreted by *Pseuduvaria mulgraveana* (yellow arrows). (f) Drosophilid fly consuming nectar in a *Pseuduvaria hylandii* flower. Photo credit: (a–f): Ming-Fai Liu.

### Floral biology and pollinator activities—*Polyalthia* species

*Polyalthia xanthocarpa* (Fig. 1f) exhibited a 2–3-day pistillate phase followed by a 1-day staminate phase (Table 1). The transition between pistillate and staminate phase was not observed directly and could have occurred at night: it is therefore unclear whether an interim or overlap phase was present. As with *Meiogyne*, *Po. xanthocarpa* bore yellow flowers with a banana-like odour (Fig. 1f) and were visited by curculionid beetles that accessed the floral sexual organs in both the staminate and pistillate phases (Table 1). Pollen grains were not observed on the limited number of pistillate-phase visiting beetles, however. The curculionid beetles were observed to consume pollen and utilise the flower for shelter. Secretion of stigmatic exudate in *Po. xanthocarpa* was weak and the consumption of stigmatic exudate was not observed. In contrast, *Polyalthia hispida* bore yellow flowers without detectable odour (Fig. 1e). Its anthesis comprised a three-day pistillate phase followed by a two-day staminate phase. No floral visitors were observed.

### Floral biology and pollinator activities—*Uvaria* species

*Uvaria concava* bore flowers with red petals and emitted a fruity odour (Fig. 1k). It had a 2– 3-day pistillate phase, followed by a prolonged overlap phase (1–2 days), in which stigmas remained attached to the ovaries as the anthers dehisce. The drying and abscission of stigmas usually occurred one day after the overlap phase and were sometimes synchronised with petal abscission. In most flower, petal abscission occurred two days after the staminate function initiated. Because of the variation in the timing of stigma drying, the duration of staminate phase varied between 0–1 day. Nitidulid beetles (subfamily Nitidulinae) were the primary floral visitors observed: they accessed the floral sexual organs and were present throughout anthesis. They were probable pollinators since pollen-laden individuals were found in pistillate-phase flowers. We observed the visitation of a single honeybee (*Apis* sp.) during the overlap phase; although the bee contacted the floral sexual organs while collecting pollen, we failed to capture it and were unable to assess pollen deposition.

*Uvaria rufa* flowers bore orange petals with tints of red and green (Fig. 1l). The flower emitted a fruity odour and had a one-day pistillate phase, followed by a one-day interim phase and a four-day staminate phase. Only one unidentified fly was observed, which visited during the interim phase. It is unclear whether pollen grains were deposited onto this fly visitor.

### Floral biology and pollinator activities—*Pseuduvaria*

*Pseuduvaria* species (Fig. 1g–j) displayed an array of sexual systems (Table 1). *Pseuduvaria hylandii* was assessed to be dioecious, while *Ps. mulgraveana* and *Ps. glabrescens* were assessed to be structurally andromonoecious, in which the same individuals bear staminate flowers that lack carpels, as well as bisexual flowers with a few stamens. *Pseuduvaria hylandii* had prolonged anthesis for both pistillate and staminate flowers (> 6 days), but we were unable to assess the anthesis duration (and floral phenology) for *Ps. glabrescens* due to logistic limitations. Anthesis in staminate flowers of *Ps. mulgraveana* lasts two days, whereas in hermaphroditic flowers the pistillate phase lasts for two days, followed immediately by petal abscission, with anthers dehiscing two days after petal abscission. We were only able to assess one staminate individual of *Ps. villosa* and were therefore unable to determine its sexual system.

The flowers of all four *Pseuduvaria* species possessed white outer petals (Fig. 1g–j). They also possessed white to reddish inner petals with dark maroon inner petal glands on their adaxial surface (Fig. 2d). These flowers emitted a strong fermentation odour that resembled rotten fruits. They had a mitriform corolla, with large basal apertures that allowed larger insects to access the floral chamber. The inner petals were apically connivent and the three pairs of glands on the inner petals form a platform in the floral chamber (Fig. 1h, Fig. 2d–f). Nectar secretion was observed on the inner petal glands and surrounding dark red petal tissues of the four *Pseuduvaria* species (Fig. 2e). Drosophilid flies and nitidulid beetles were observed to be the primary floral visitors of *Ps. glabrescens*, *Ps. hylandii* and *Ps. mulgraveana* (Table 1). These insects were observed accessing the floral chamber and consuming nectar, often positioning themselves underneath the sexual organs to access the nectary. Pollen grains were observed to be deposited either onto the back of the insects (Fig. 2f), or onto the inner petal glands where the pollen grains were then picked up secondarily by the flies and beetles on their legs. The drosophilid flies and nitidulid beetles were observed moving across the stigmas and stamens with their legs touching the sexual organs, which may assist pollen transfer. Some drosophilid flies are large enough to brush the back of their thorax or abdomen against the sexual organs, which may additionally deposit pollen grains onto the stigmas. Both drosophilid flies and nitidulid beetles were observed to access the sexual organs of pistillate and staminate flowers for *Ps. glabrescens*, *Ps. hylandii* and *Ps. mulgraveana*. While pollen grains were observed on the drosophilid flies and nitidulid beetles found in pistillate-phase flowers of *Ps. hylandii*, pollen deposition was observed only on nitidulid beetles from pistillate-phase flowers of *Ps. glabrescens* and *Ps. mulgraveana*. Other dipteran visitors, namely lauxaniid and neriid flies, were also observed in *Ps. hylandii*, but they are too large to access the floral chamber and thus unlikely to be pollinators. Unfortunately, we failed to observe any floral visitors for *Ps. villosa*, though it has similar floral colour and odour to the other *Pseuduvaria* species (Fig. 1j).

### Floral biology and pollinator activities—*Monoon*

*Monoon patinatum* produced flowers with greenish petals (Fig. 1d). Only two staphylinid beetle visitors from the subfamily Tachyporinae were observed during the pistillate phase (Table 1). Pollen grains were not observed on their bodies.

### Floral Colour

Various types of floral coloration were observed in the studied species, including red (*Uvaria concava*), orange (*U. rufa*), yellow (*Meiogyne cylindrocarpa*, *Me. trichocarpa*, *Me. stenopetala*, *Polyalthia hispida* and *Po. xanthocarpa*), green (*Monoon patinatum*), and a mixture of white and maroon (*Pseuduvaria hylandii*, *Ps. glabrescens*, *Ps. mulgraveana* and *Ps. villosa*). The spectral reflectance of the petals of 12 Annonaceae species is shown in Fig. 3. The petals of all species are not UV-reflective from 300–350 nm; most of the reflectance was observed in the visible spectrum (400–700 nm). Note that while yellow-flowered species possess similar coloration to human perception, their spectral reflectance can be quite varied, highlighting the importance of quantification of reflective spectra.

**Fig. 3.**
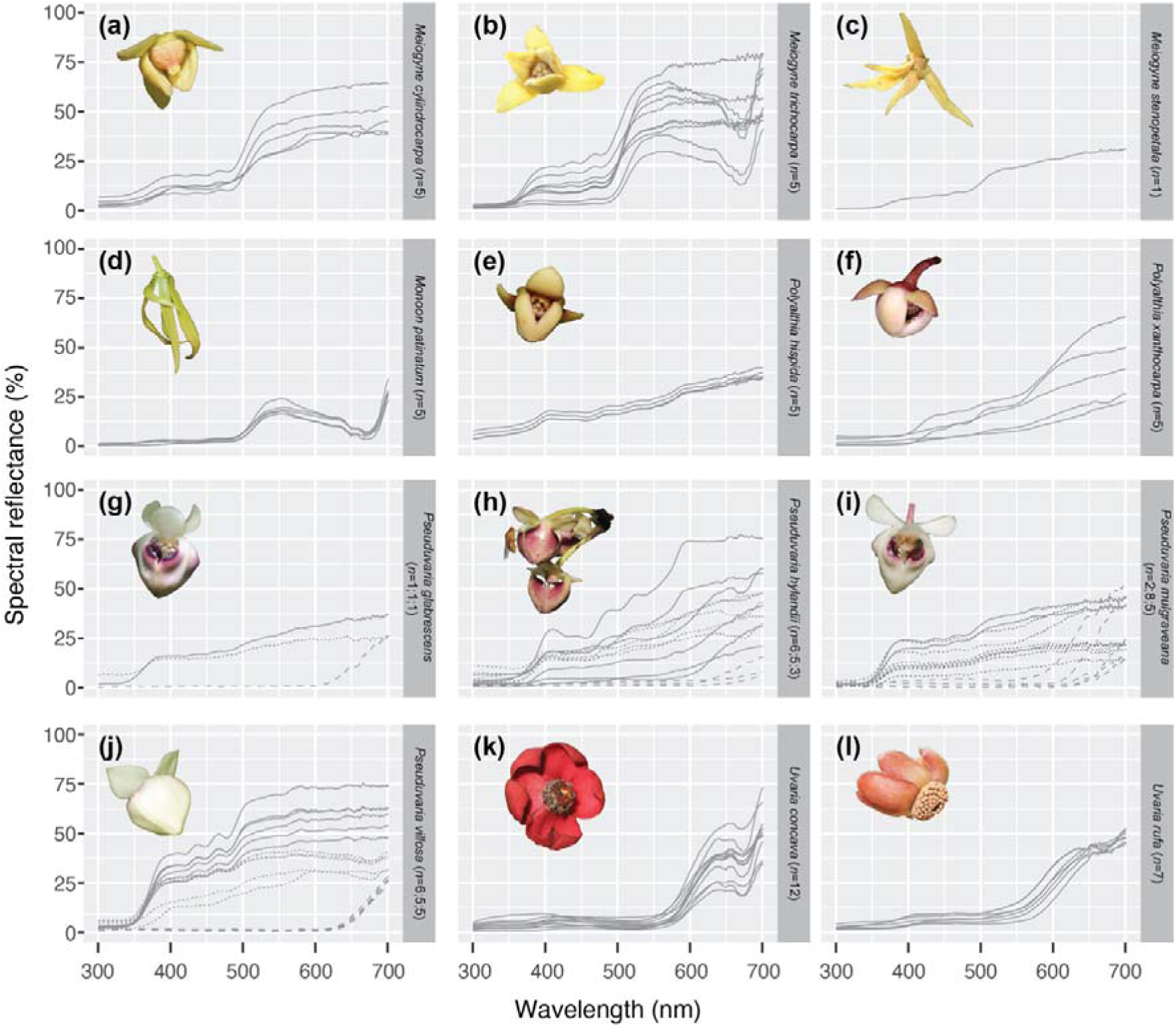
Spectral reflectance of 12 Annonaceae species. (a) *Meiogyne cylindrocarpa*. (b) *Meiogyne trichocarpa*. (c) *Meiogyne stenopetala*. (d) *Monoon patinatum*. (e) *Polyalthia hispida*. (f) *Polyalthia xanthocarpa*. (g) *Pseuduvaria glabrescens*. (h) *Pseuduvaria hylandii*. (i) *Pseuduvaria mulgraveana*. (j) *Pseuduvaria villosa*. (k) *Uvaria concava*. (l) *Uvaria rufa*. For species with drooping inner petals and no colour differentiation between outer and inner petal whorls, the inner petal abaxial surface was measured (a–f; i.e. *Meiogyne*, *Monoon* and *Polyalthia*). For species with spreading inner petals and no colour differentiation between outer and inner petal whorls, the inner petal adaxial surface (k–l; i.e. *Uvaria*) was measured. For species with dissimilar petal whorls (g–j; i.e. *Pseuduvaria*), three perianth areas were measured (solid line: inner petal abaxial surface; dotted line: outer petal adaxial surface; dashed line: inner petal gland); the sample size (*n*) for these species refer to the sample size for these three perianth areas in the aforementioned order. Photo credits: (a, b, d, e, g–l): Ming-Fai Liu; (c): Chun-Chiu Pang; (f): Garry & Nada Sankowsky.

### Floral Odour

The floral headspace composition of 11 Annonaceae species is summarised in Table 2. We detected a total of 97 volatiles from seven classes of compounds. We did not detect any major volatiles in the headspace of *Polyalthia hispida* against the ambient air control, and therefore did not present the headspace data here.

**Table 2.**
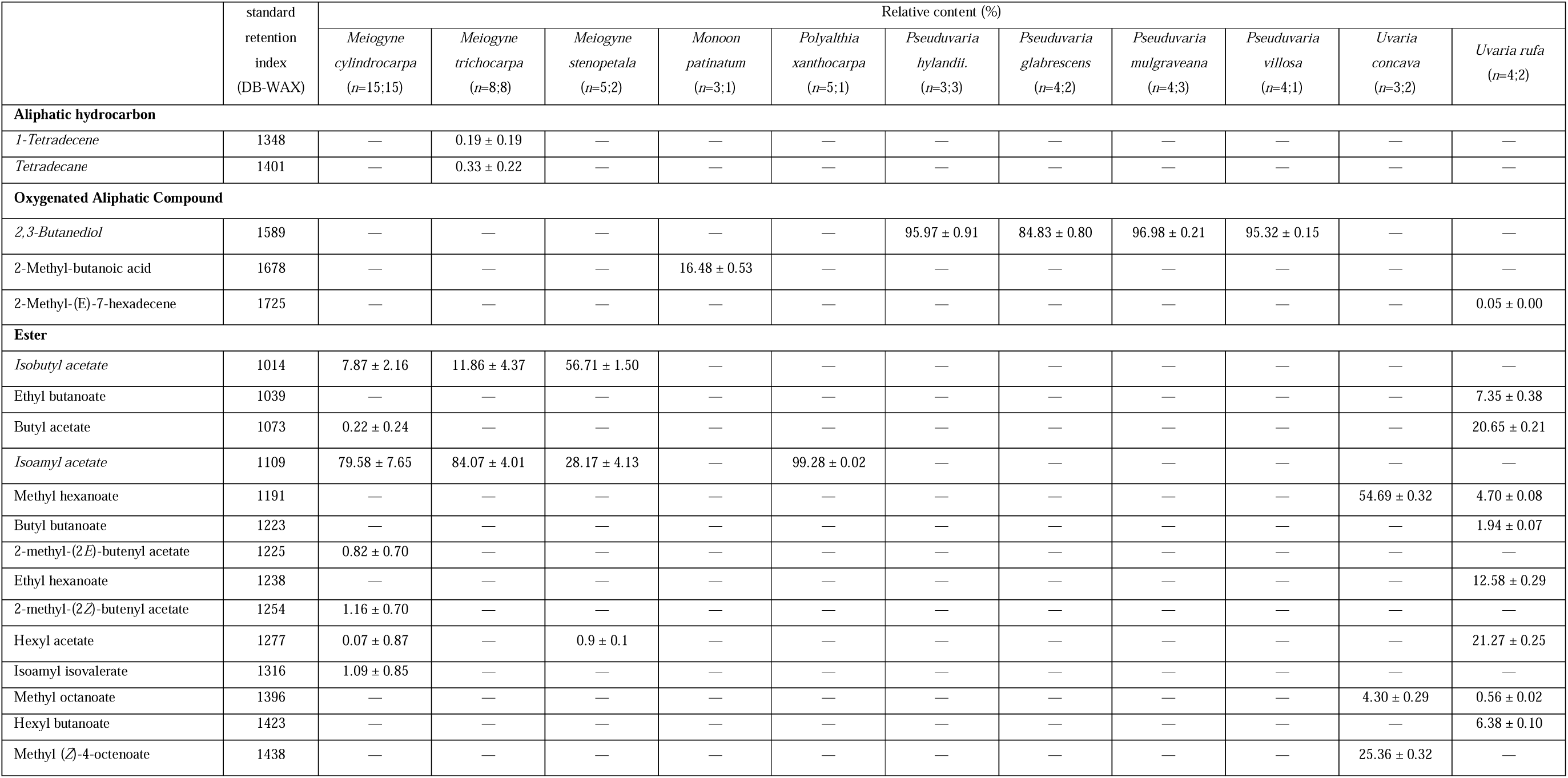
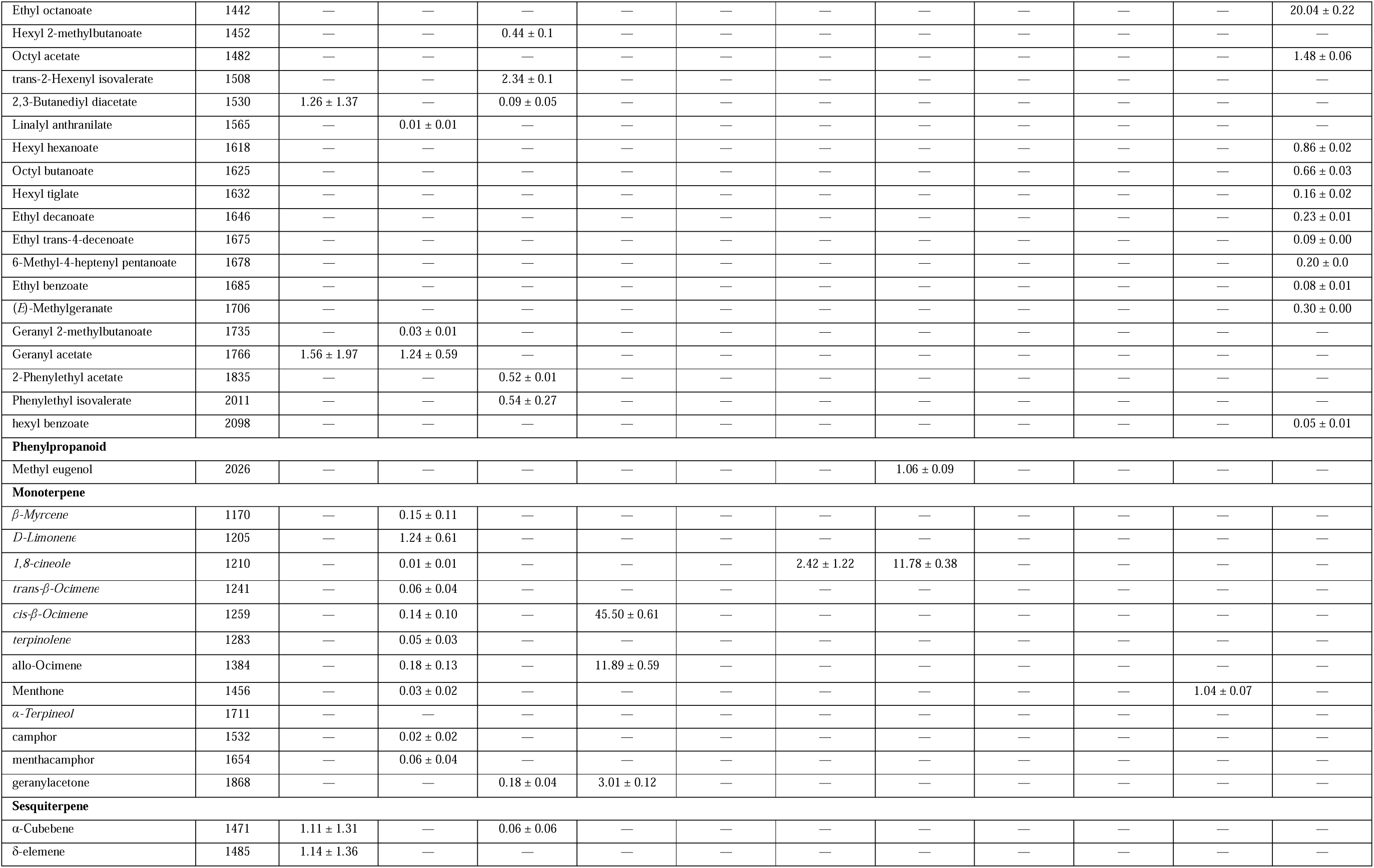
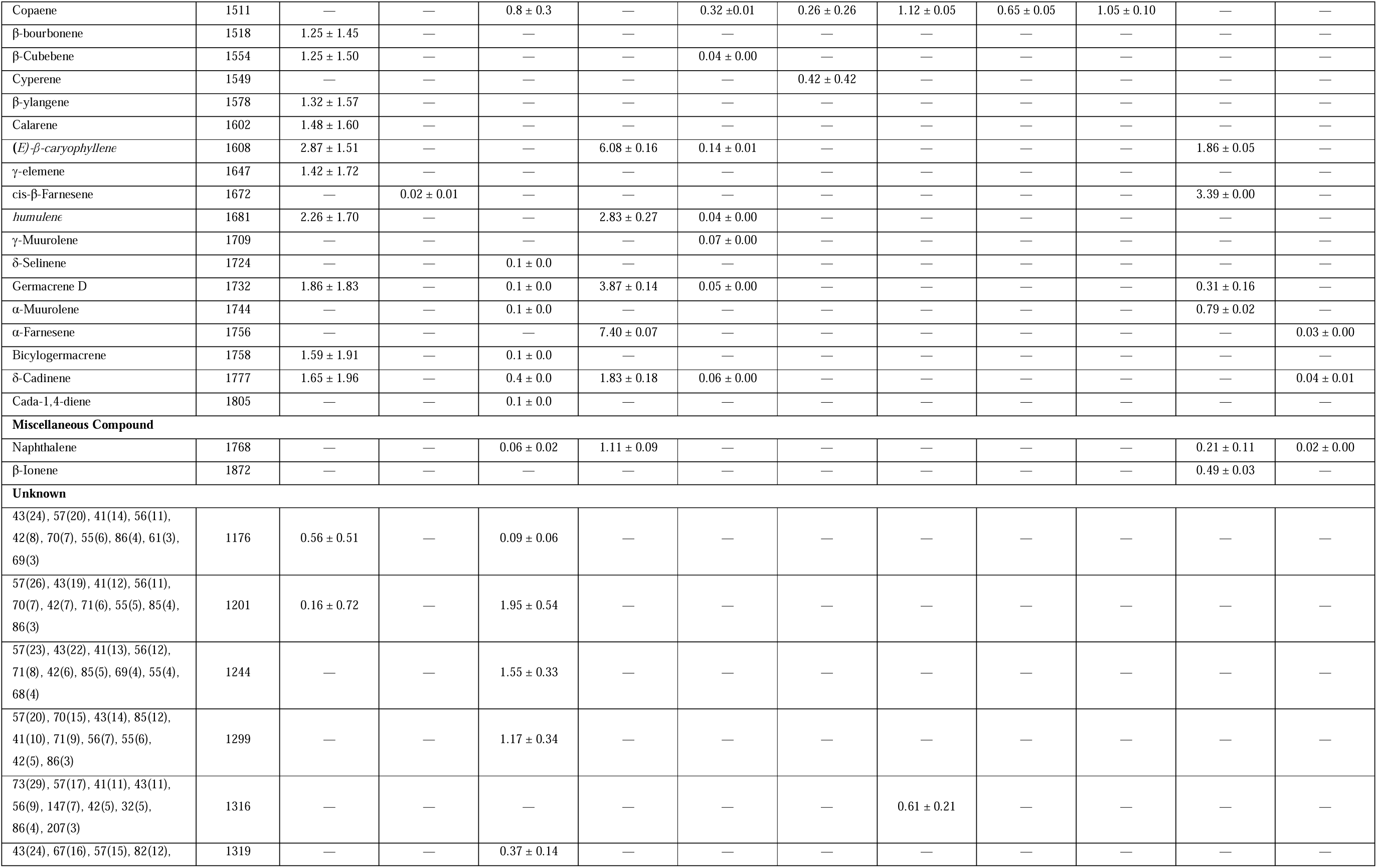
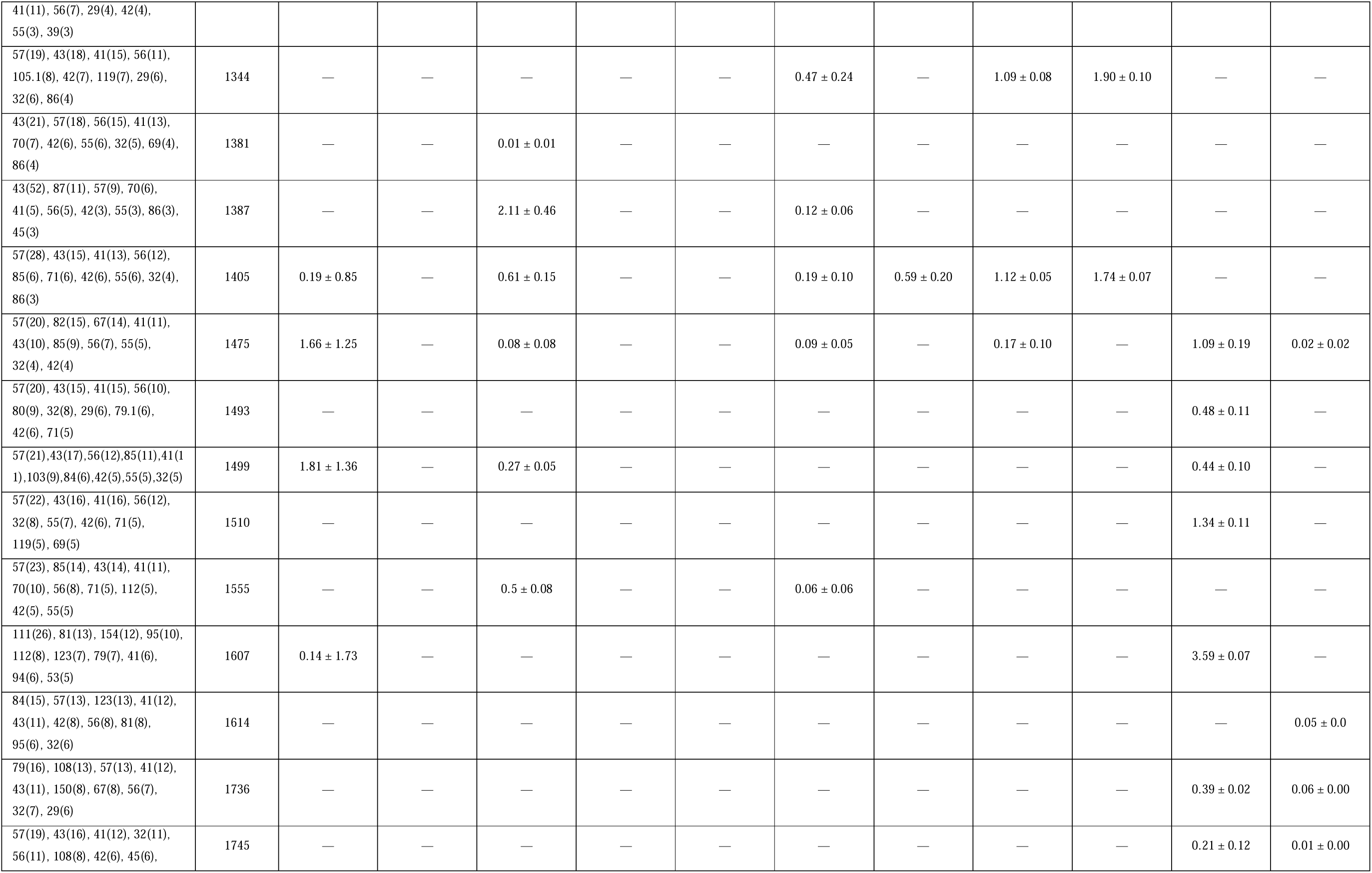
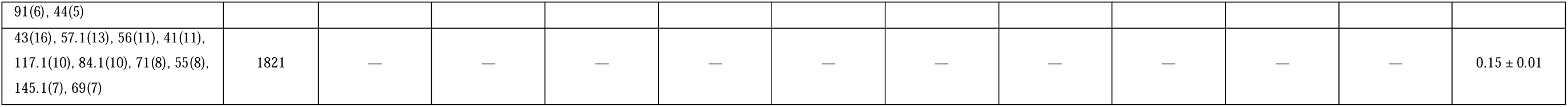
Floral scent composition of *Meiogyne*, *Monoon*, *Polyalthia*, *Pseuduvaria* and *Uvaria* species. The average percentage of the relative peak area for each volatile and its standard error are presented below. Italicised molecules were verified by co-injecting with commercially available standards. Unknown molecules were denoted by the top ten most abundant ion fragments, followed by their relative intensity in parentheses. *n* refers to the number of technical replicates performed, followed by the number of individuals the odour was collected from.

Aliphatic ester was the most abundant class of volatiles in *Meiogyne cylindrocarpa*, *Me*. *trichocarpa*, *Me*. *stenopetala*, *Polyalthia xanthocarpa*, *Uvaria concava* and *U*. *rufa*. The headspaces of all *Meiogyne* species and *Po*. *xanthocarpa* contained a varying degree of esters, including isobutyl acetate (*Me*. *cylindrocarpa*: 7.87 ± 2.16%; *Me*. *trichocarpa*: 11.86 ± 4.37%; *Me*. *stenopetala*: 56.71 ± 1.5%) and isoamyl acetate (*Me*. *cylindrocarpa*: 79.58 ± 7.65%; *Me*. *trichocarpa*: 84.07 ± 4.01%; *Me*. *stenopetala*: 28.17 ± 4.13%; *Po*. *xanthocarpa*: 99.28 ± 0.02%). For *U*. *concava*, the floral headspaces were dominated by two esters, namely methyl-(*Z*)-4-octenoate (25.36 ± 0.32%), and methyl hexanoate (54.69 ± 0.32%), while for *U*. *rufa*, the most abundant volatiles were hexyl acetate (21.27 ± 0.25%), butyl acetate (20.65 ± 0.21%), and ethyl octanoate (20.04 ± 0.22%). In *Meiogyne* and *Polyalthia*, branched-chain aliphatic esters dominated the floral odour, while in *Uvaria*, the floral odours primarily consisted of straight-chain the aliphatic esters. For *Monoon patinatum*, the headspace mainly comprised the monoterpenes β-ocimene (45.50 ± 0.61%) and allo-ocimene (11.89 ± 0.59%), and 2-methyl-butanoic acid (16.48 ± 0.53%). The oxygenated aliphatic compound was the most abundant volatile class for *Pseuduvaria hylandii*, *Ps*. *glabrescens*, *Ps*. *mulgraveana* and *Ps*. *villosa*, consisting of the molecule 2,3-butanediol (84.83–96.98% of the headspaces).

The Annonaceae floral odours detected in this study and those derived from the literature were visualised in an NMDS plot (Fig. 4), based on 315 volatiles from 40 species, in which 11 was newly generated in this study. The 2D stress level of the NMDS was 0.140. The global ANOSIM statistic R based on the type of pollination type was 0.4528, at *p*-value of 0.00001***, suggesting floral scents are significantly correlated to the type of pollination system. The floral odours of small-beetle pollinated species in the literature cluster with the small-beetle pollinated species identified in this study, including *Meiogyne cylindrocarpa*, *Me. trichocarpa*, *Me. stenopetala*, and *Po. xanthocarpa*. The species with unknown pollinator, *Monoon patinatum*, was also retrieved nested inside this clusters. Similarly, the *Pseuduvaria* species pollinated by flies and beetles identified in the current study occupied a similar space with *Asimina* species that are pollinated by flies and beetles, suggesting there is a convergence in floral odour for species pollinated by a mixture of flies and beetles. The bee- and beetle-pollinated species *Uvaria concava* did not cluster with other bee-pollinated Annonaceae species, but instead occupied a distinct olfactory space.

**Fig. 4.**
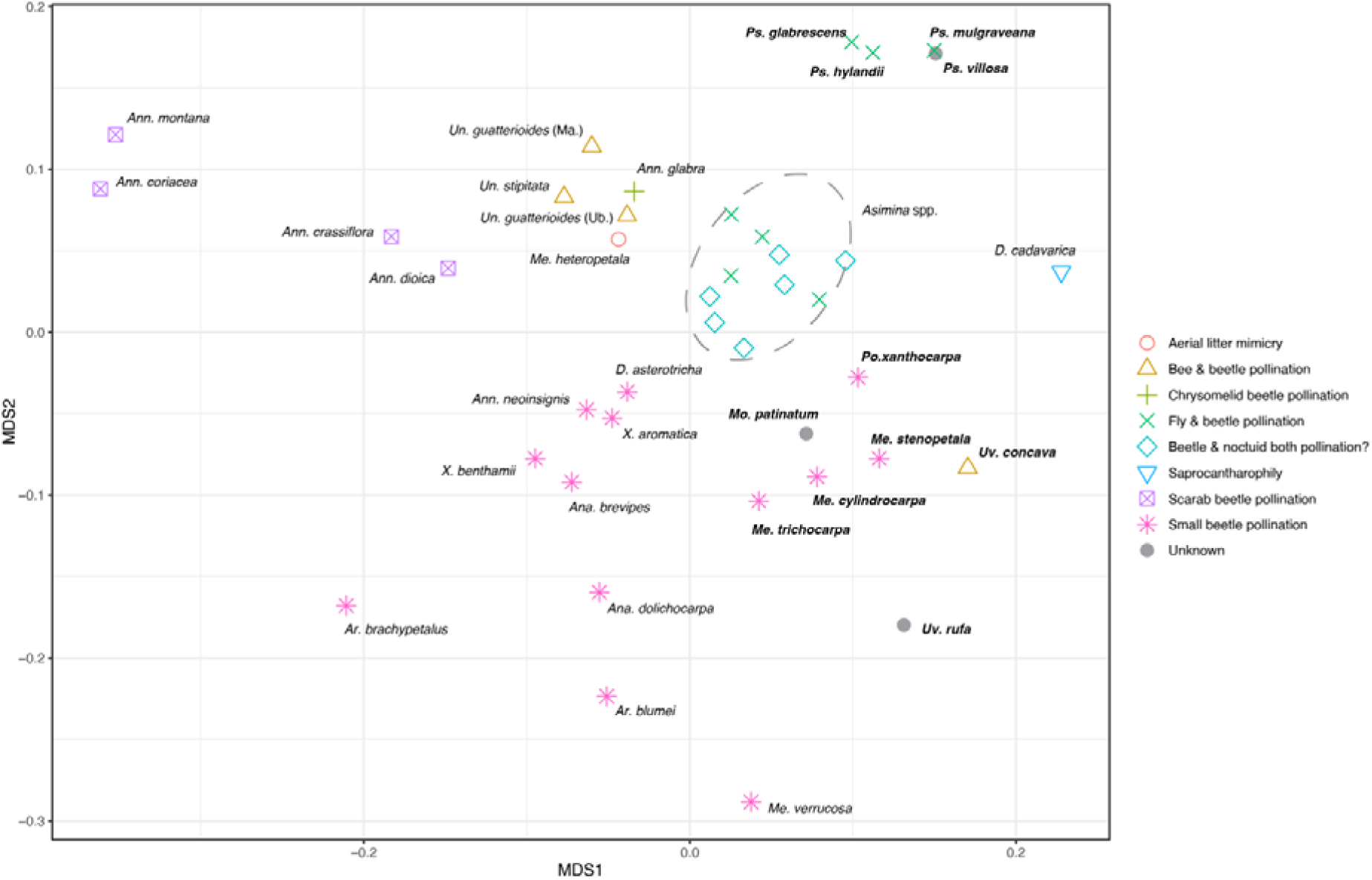
Non-metric dimensional scaling (NMDS) plot of Annonaceae floral scents constructed based on Bray-Curtis dissimilarity index. Species with newly obtained floral odour was indicated by bold font. (2D stress: 0.140; global ANOSIM statistic R: 0.4528, *p*-value: 0.0001***) (*Ana*. *brevipes*: *Anaxagorea brevipes*; *Ana*. *dolichocarpa*: *Anaxagorea dolichocarpa*; *Ann*. *montana*: *Annona montana*; *Ann*. *coriacea: Annona coriacea*; *Ann*. *crassiflora*: *Annona crassiflora*; *Ann*. *dioica*: *Annona dioica*; *Ann*. *neoinsignis*: *Annona neoinsignis*; *Ann.* glabra: *Annona glabra*; *Ar. blumei*: *Artabotrys blumei*; *Ar. brachypetalus*: *Artabotrys brachypetalus*; *D*. *asterotricha*: *Duguetia asterotricha*; *D*. *cadavarica*: *Duguetia cadavarica*; *Me*. *cylindrocarpa*: *Meiogyne cylindrocarpa*; *Me*. *trichocarpa*: *Meiogyne trichocarpa*; *Me*. *stenopetala*: *Meiogyne stenopetala*; *Me. verrucosa*: *Meiogyne verrucosa*; *Mo. patinatum*: *Monoon patinatum*; *Po. xanthocarpa*: *Polyalthia xanthocarpa*; *Ps. glabrescens*: *Pseuduvaria glabrescens*; *Ps. hylandii*: *Pseuduvaria hylandii*; *Ps. mulgraveana*: *Pseuduvaria mulgraveana*; *Ps. villosa*: *Pseuduvaria villosa*; *Un. guatterioides* (Ma.): *Unonopsis guatterioides* (Manaus); *Un. guatterioides* (Ub.): *Unonopsis guatterioides* (Uberlandia); *Un. stipitata*: *Unonopsis stipitata*; *Uv. concava*: *Uvaria concava*; *Uv. rufa*: *Uvaria rufa*; *X. aromatica*: *Xylopia aromatica*; *X. benthamii*: *Xylopia benthamii*.

## DISCUSSION

### Floral phenology

Most species assessed in this study exhibited a 3–5 day anthesis period and none exhibited circadian trapping, a time-dependent trapping mechanism in which pollinators are trapped and released in accordance to the insects’ peak circadian activities (Lau et al., 2017). The duration of anthesis for most species in the current study is similar to the typical non-trapping Annonaceae species (≥ 2 days) (Pang and Saunders, 2014; Lau et al., 2017). As the family is self-compatible (Pang and Saunders, 2014), temporal separation of sexual phases is an important adaptation to reduce autogamy. An interim phase was observed in *Uvaria rufa*, during which the stigmas are no longer receptive and anther dehiscence has not begun, potentially offering additional assurance to prevent autogamy (Pang and Saunders, 2014). However, in some of the other species, including *Meiogyne* spp. and *U*. *concava*, an overlap phase was observed, during which carpels and stamens are simultaneously functional within a flower. This overlap phase could offer reproductive assurance by autogamy when pollinator availability is scarce. It was previously reported that *U*. *concava* was pollinated by stingless bees (Meliponinae) in its natural population (50 km from our site of study) (Silberbauer-Gottsberger et al., 2003). Because bee visitors to Annonaceae often collect pollen only as a food reward, and seldom visit pistillate-phase flowers, the overlap phase has been suggested to be essential for bee pollination (Gottsberger, 2014). It was, however, reported that meliponine visitors also collect stigmatic exudate as a reward (Silberbauer-Gottsberger et al., 2013), suggesting the flower offers reward to the meliponine visitors even during pistillate phase. The role of the prolonged overlap phase for *U*. *concava* is therefore contentious.

Floral phenology also played a role in the sexual function of the flower, especially for *Pseuduvaria* species. Previous studies have shown that *Ps*. *mulgraveana* is structurally andromonoecious (Pang et al., 2013), meaning that staminate flowers and bisexual flowers are borne on the same plant. The stamens of bisexual flowers, however, release functional pollen grains only after petal abscission (Pang et al., 2013), rendering the plant functionally monoecious, which is corroborated by our observations.

### Inner petal corrugation and glands

Inner petal outgrowth was observed in *Meiogyne* spp. and *Pseuduvaria* spp. in the current study. This outgrowth has independently evolved in around 20 genera, including *Annona* L., *Alphonsea* Hook.f. & Thomson, Asimina Adans., *Asteranthe* Engl. & Diels, *Duguetia* A.St.-Hil., *Maasia* Mols, Kessler & Rogstad, *Meiogyne* Miq., Miliusa Lesch. ex A.DC., *Orophea* Blume, *Porcelia* Ruiz & Pav., *Pseuduvaria* Miq., *Sapranthus* Seem., *Stenanona* Standl., *Tetrameranthus* R.E.Fr., *Tridimeris* Baill., *Uvaria* L., *Wangia* X.Guo & R.M.K.Saunders, and *Xylopia* L. (van Heusden, 1992; Schatz and Maas, 2010; Xue et al., 2017). In *Pseuduvaria*, the inner petal outgrowth has been reported to be a nectary gland (Silberbauer-Gottsberger et al., 2003). In the current study, the four *Pseuduvaria* species also possessed this floral tissue, which invariably functioned as nectary, corroborating the previous study. Likewise, all three *Meiogyne* species assessed in our study possessed weakly folded to longitudinally grooved basal growths on the adaxial surface of the inner petals. Although this structure has previously been interpreted as a gland (van Heusden, 1992, 1994), histological studies have failed to identify secretory ducts or hairs (Xue et al., 2021). The inner petal corrugation may potentially offer tactile cues for the floral mimicry of aerial litter in *Meiogyne heteropetala* (Liu et al., 2024), but the role of this tissue remains enigmatic for most *Meiogyne* species. Our observations indicate that pollinators preferentially consume the inner petals over other floral structures (Fig. 2), suggesting that the corrugation likely functions as a food body in *Me*. *cylindrocarpa*, *Me*. *trichocarpa* and *Me*. *stenopetala*. In Annonaceae, specialised nutritious petal tissues are common in species that are pollinated by scarab beetles in the neotropics, including *Annona*, *Cymbopetalum*, *Duguetia*, and *Malmea* species (Gottsberger and Webber, 2018). Among these lineages with specialised food body, there is potential convergence in tissue structure. The inner petal corrugation of these species is anatomically similar to those found in *Meiogyne*, lacking secretory openings, ducts or trichromes, and comprising a mixture of cells full of starch granules and cells enriched with tannins (Gottsberger and Webber, 2018; Xue et al., 2021).

### Exploitation of pollinators associated with fruits

The fruity floral aromas and beetle pollination of Annonaceae have led some authors to propose some Annonaceae flowers mimic fruits (Johnson and Schiestl, 2016; Goodrich and Jürgens, 2018). In particular, fruit mimicry has been postulated in Anaxagorea, Annona, Artabotrys, Duguetia and Xylopia and is likely to be prevalent in the Annonaceae (Johnson and Schiestl, 2016; Goodrich and Jürgens, 2018; Chen et al., 2020). These flowers are pollinated by small beetles such as Curculionidae, Nitidulidae and Staphylinidae that are known to use fruits as a food source or oviposition site (Phelan and Lin, 1991; Racette et al., 1992; Frank and Thomas, 1999; Mutinelli et al., 2015).

Small beetle pollinators (families Nitidulidae, Curculionidae and Staphylinidae) were observed as floral visitor of *Me*. *cylindrocarpa*, *Me*. *trichocarpa*, *Me*. *stenopetala* and *Po*. *xanthocarpa* (Table 1), which are largely congruent with existing literature. Previous studies have recorded small beetle visitors in *Polyalthia xanthocarpa* flowers (as “*Haplostichanthus* spec. (“Cape tribulation”)”; Morawetz, 1988), and curculionid beetles have been observed visiting the flowers in Cape Tribulation, Australia (pers. comm., Dr David Tng). For *Meiogyne*, nitidulid and curculionid beetles have been reported to be the pollinators of *Me*. *cylindrocarpa* in tropical rainforests in West Malesia (Momose, 2005; Roubik et al, 2005). Beetles from these families were also observed in the current study.

The spectral reflectance of *Meiogyne* species, especially *Me*. *cylindrocarpa* and *Me*. *trichocarpa*, exhibited low UV reflectance, with stronger reflectance from 500–700 nm (Fig 3). The UV-absorbing yellow spectral profile is a characteristic for pollen (Dötterl et al., 2014) and was also reported in fruits such as bananas (Xie et al., 2018). Some nitidulid and curculionid beetles are known to associate with fruits (Phelan and Lin, 1991; Racette et al., 1992; Mutinelli et al., 2015), and nitidulidae has previously been demonstrated to be attracted to various shade of yellow (Döring et al., 2012; Vuts et al., 2022). Floral aromas resembling ripe fruits were recorded in *Me*. *cylindrocarpa*, *Me*. *trichocarpa*, *Me*. *stenopetala*, *Po*. *xanthocarpa*. Their floral odour resembled strong banana-like scent to human perception, and is largely composed of either isoamyl acetate or isobutyl acetate (Table 2). These two molecules have been reported in various fruits, including banana (Jordán et al., 2001), papaya (Katague and Kirch, 1965), pear (Zhang et al., 2023) and wild soursop (Pino et al., 2002), and is implicated as the semiochemicals of Curculionidae and Nitidulidae (Rochat et al., 2000; Torto et al., 2007). The NMDS revealed that all these small-beetle pollinated species clustered with other Annonaceae species that are also pollinated by small insects, such as nitidulid, curculionid and staphylinid beetles (Fig. 4; Jürgens, 2000; Chen et al., 2020). These flowers also emit floral odour largely composed of aliphatic esters (Jürgens, 2000; Chen et al., 2020), suggesting there might be a specialisation in floral odour for Annonaceae species that utilise this guild of pollinators. The current evidence suggests that *Me*. *cylindrocarpa*, *Me*. *trichocarpa*, *Me*. *stenopetala* and *Po*. *xanthocarpa* likely attract their pollinators at least by exploiting visual and olfactory cues that indicate fruits.

### Exploitation of pollinators associated with fermented substrates

Records of floral visitors to *Pseuduvaria* are more comprehensive than for most other Annonaceae genera in Australia. In a previous pollination study on another Australian *Pseuduvaria* species, *Ps*. *froggattii*, it was shown that the flowers are primarily pollinated by drosophilid flies (Silberbauer-Gottsberger et al., 2003). Similarly, *Ps*. *glabrescens* was previously reported to be visited primarily by small flies (as *Pseuduvaria* spec. (“Davies Creek”); Morawetz, 1988). *Pseuduvaria mulgraveana*, on the other hand, was reported to be pollinated by the nitidulid beetle, *Aethina australis*, and was only infrequently visited by flies. Our observations are largely congruent with these studies: *Ps*. *hylandii*, *Ps*. *glabrescens* and *Ps*. *mulgraveana* were observed to be visited by both nitidulid beetles and drosophilid flies. Pollen deposition on the nitidulid beetles was observed in the pistillate-phase flowers of all three species, while drosophilid flies appeared to be the pollinator only for *Ps*. *hylandii*. It was previously postulated that flies could pollinate *Ps*. *villosa*, but floral visitors were not observed in our study.

Floral coloration in the *Pseuduvaria* species varies across different areas of petal tissues (Figs. 1 & 3). The abaxial surface of the inner petals in most species demonstrate largely similar reflectance spectra with the adaxial surface of the outer petals, except for *Ps*. *hylandii*. In contrast to these areas, the inner petal glands of all four *Pseuduvaria* species studied are burgundy red with generally lower reflectance across all wavelengths. Additionally, all four *Pseuduvaria* species emit an odour reminiscent of rotten fruits. Rotting fruits have been reported to display a reduction in overall reflectance across the visible spectrum under fungal infection (Liu et al., 2020). The contrast between the dark burgundy inner petal gland and the remaining bright floral tissues may have evolved to develop similar visual cues as rotten patches on fruits. Interestingly, in *Pseuduvaria*, nectar secretion is localised in the darkened area (Fig. 2), suggesting there is co-localisation of gustatory cue and visual cue, which may help position the pollinators underneath the reproductive organs and facilitate pollen transfer. The floral odours of all four *Pseuduvaria* species are largely composed of 2,3-butanediol (Table 2), which is a common volatile emitted by yeast and fermenting substrates (Goodrich et al., 2006, 2023). This molecule is one of the major volatiles of *Asimina triloba* (Goodrich et al., 2006), a maroon-petalled floral mimic of fermentation substrates (Goodrich et al., 2023). As with *Ps*. *glabrescens*, *Ps*. *hylandii* and *Ps*. *mulgraveana*, *Asimina triloba* is pollinated by small flies and small beetles (Martin, 2021; Goodrich et al., 2023). This suggests convergence in floral colour and odour between *Asimina* and *Pseuduvaria* for attracting the same pollinator guilds, with the NMDS showing that these taxa share a similar olfactory space (Fig. 4). The four *Pseuduvaria* species may therefore be potential candidates for floral mimics of fermentation substrates.

### Exploitation of bee and beetle pollinators

*Uvaria concava* was reported to be pollinated by stingless bees in the wild (Silberbauer-Gottsberger et al., 2003) but is here inferred to be pollinated by nitidulid beetles in cultivation (Table 1). However, the cultivated environment is only *ca.* 50 km from the wild population where the previous study was conducted. This species possesses bright-red petals, reflecting the visible spectrum mostly at 600–700 nm (Fig. 3). However, neither the tribe Meliponini (stingless bees) nor the family Apidae have photoreceptors within this range (van der Kooi et al., 2021). Though the photosensitivity of Nitidulidae is unknown, most coleopteran families with known visual systems are unable to detect this part of the visible spectrum (van der Kooi et al., 2021). It is therefore unclear how floral visual cues assist pollinator attraction in *U*. *concava*. The floral odour of *U*. *concava* is largely composed of methyl hexanoate and methyl (*Z*)-4-octenoate. While there is little documentation how meliponine bees and nitidulid beetles respond to these molecules, methyl hexanoate is reported to be a pheromone for male bumblebees (Valterová et al., 2001). Among Annonaceae species that emit esters, branched-chain aliphatic esters are typically produced (Table 2; Jürgens et al, 2000), but *Uvaria* spp. primarily produce straight-chain aliphatic esters instead. Interestingly, the branched-chain aliphatic ester isoamyl acetate (the dominant floral scent component of *Meiogyne* spp. and *Polyalthia xanthocarpa*) is a bee alarm pheromone that reduces foraging in the honeybee species *Apis mellifera* and *Apis cerana* (Gong et al., 2017). Nonetheless, whether this structural difference in floral esters contributes to the shift to bee pollination remains to be investigated. Interestingly, *U. concava* occupied a distinct locus in the odour space from other bee-pollinated Annonaceae species, but instead is more closely positioned with small-beetle pollinated Annonaceae species. It might be attributed to the different lineage of the pollinator. *Unonopsis* spp. were pollinated by XXXXXXX, instead of Apidae. Future studies using colour-based and scent-based bioassays could help assess the role played by visual and olfactory cues in bee-pollinated Annonaceae species.

## Conclusions

The basic floral biology, petal spectral reflectance and floral scent of selected species in five Annonaceae genera are characterised here in detail. Our study has identified multiple species that have floral traits resembling fruits (*Meiogyne* spp. and *Po. xanthocarpa*) and fermented substrates (*Meiogyne* spp. and *Po*. *xanthocarpa*). The specialised floral cues and adaptations identified in this study provide preliminary evidence for potential floral mimicry of fruits and fermented substrates, and offer insights into how floral cues may attract floral visitors. These findings serve as a prelude to future studies on specialised pollination systems, and provide baseline information on floral traits in the pantropical family Annonaceae. Future studies involving the characterisation of colour and odour of co-occurring fruits and fermenting substrates would be important to assess whether these species exploit their pollinators through floral mimicry.

## ACKNOWLEDGEMENTS

The authors thank Garry and Nada Sankowsky, Graham Wood, Charles Clarke (Cairns Botanic Gardens), and Janelle Jung (Gardens by the Bay) for allowing us to work on the living collections of Annonaceae species in their arboreta or botanic gardens. The authors also thank Frank Zich from Australian Tropical Herbarium, and the staff from Mount Tamborine National Park and the Queensland Department of Environment, Science and Innovation for their assistance, and Jessie Lai and Laura Wong for their technical support. This research is funded by the Hong Kong Research Grants Council (HKU17112616), awarded to R.M.K.S.

## AUTHOR CONTRIBUTION

Ming-Fai Liu, Junhao Chen and Richard M. K. Saunders planned the project. Ming-Fai Liu, Junhao Chen, Chun-Chiu Pang and Tanya Scharaschkin conducted the field work. Ming-Fai Liu conducted the lab experiments, performed the analysis and wrote the manuscript. Ming-Fai Liu, and Richard M. K. Saunders provided interpretation. All authors edited the manuscript.

## DATA AVAILABILITY STATEMENT

All data generated in this study is included in the main text of this article.

